# Arrestin recognizes GPCRs independently of the receptor state

**DOI:** 10.1101/2024.07.19.604263

**Authors:** Ivana Petrovic, Meltem Tatli, Samit Desai, Anne Grahl, Dongchun Ni, Henning Stahlberg, Anne Spang, Stephan Grzesiek, Layara Akemi Abiko

**Author notes:** Correspondence (S.G), (L.A.A).

## Abstract

Only two non-visual arrestins recognize many hundreds of different, intracellularly phosphorylated G protein-coupled receptors (GPCRs). Due to the highly dynamic nature of GPCR•arrestin complexes, the critical determinants of GPCR-arrestin recognition have remained largely unclear. We show here that arrestin2 recruitment to the β_1_-adrenergic receptor (β_1_AR) can be induced by an arrestin-activating phosphopeptide that is not covalently linked to the receptor and that the recruitment is independent of the presence and type of the orthosteric receptor ligand. Apparently, the arrestin-receptor interaction is driven by the conformational switch within arrestin induced by the phosphopeptide, whereas the electrostatic attraction towards the receptor phosphosites may only play an auxiliary role. Extensive NMR observations show that in contrast to previous static GPCR•arrestin complex structures, the β_1_AR complex with the beta-blocker carvedilol and arrestin2 is in a G protein-inactive conformation. The insensitivity to the specific receptor conformation provides a rationale for arrestin’s promiscuous recognition of GPCRs and explains the arrestin-biased agonism of carvedilol, which largely blocks G protein binding, while still enabling arrestin engagement.

**Significance statement:** G protein-coupled receptors regulate cellular signaling through G proteins and arrestins. While G protein interactions are well understood, the molecular basis of arrestin recognition remains unclear due to the dynamic nature of GPCR•arrestin complexes. We show that arrestin recognition of the β_1_-adrenergic receptor occurs independently of the receptor’s conformational state or ligand binding. Using NMR, cryo-EM, and biochemical assays, we find that arrestin engagement is driven by a conformational change within arrestin itself, triggered by a non-receptor-attached phosphopeptide, whereas electrostatic attraction towards receptor phosphosites may only play an auxiliary role. These findings provide new insights into arrestin activation, explain its ability to recognize diverse GPCRs, with significant implications for understanding biased signaling mechanisms and designing of selective therapeutic strategies.

## Introduction

G protein-coupled receptors (GPCRs) regulate vital physiological processes by detecting a multitude of orthosteric ligands, which modulate their conformational landscape and control the signaling to intracellular effectors (1). In addition to the canonical G protein-mediated signaling (2), GPCRs can also signal via arrestin proteins (3). Hundreds of non-visual GPCRs are regulated by only two non-visual arrestin isoforms, arrestin2 and arrestin3. The binding of these arrestins to GPCRs requires phosphorylation of the receptor’s C-terminal tail or intracellular loops by GPCR kinases (GRKs) (4). This interaction can then initiate various signaling pathways independent of G-protein signaling, such as MAP kinase activation (5).

In contrast to complexes with G proteins, GPCR-arrestin complexes are highly dynamic due to the weaker nature of the receptor-arrestin interactions. As a consequence, the structures of only very few distinct GPCR-arrestin complexes have been solved so far (6–14). In all these complexes, the receptor adopts a conformation similar to the G protein-active state, characterized by a large outward displacement of the intracellular portions of transmembrane helices (TM) 5 and 6 from the helix bundle, relative to the inactive conformation. Moreover, most of these complexes required additional receptor engineering to stabilize the coupling to arrestin. Typically, this is achieved by artificially fusing the highly phosphorylated C-terminal tail of the vasopressin receptor 2 (V2Rpp) (8, 14) to the receptor of interest (9, 11, 15). V2Rpp and other phosphorylated receptor C-terminal peptides form well-defined complexes with arrestins, where the arrestin β20 strand is released from its core and replaced by the phosphopeptide forming an intermolecular β-sheet (16, 17). This interaction leads to a moderate reorientation of arrestin’s N- and C-domains and core loops. However, the forces driving the interaction with the receptor have remained unclear.

Here, we have elucidated the requirements for recruitment of arrestin2 to the β_1_-adrenergic receptor (β_1_AR) by biochemical, cryo-EM, and detailed NMR observations of both receptor and arrestin2. The only requirement for the engagement of arrestin2 with the β_1_AR is the binding of a non-receptor-attached GPCR phosphopeptide to arrestin2, with the receptor conformation and orthosteric ligand proving irrelevant. This insight explains the ability of arrestins to recognize a broad range of GPCRs as well as the mechanism of the arrestin-biased agonism of carvedilol, which is a widely used beta-blocker targeting β_1_- and β_2_-adrenergic receptors (18).

## Results

### β_1_AR forms a stable complex with arrestin2 in the presence of unligated V2Rpp, independently of orthosteric ligands

While the conformational changes within arrestin triggered by the binding of receptor phosphopeptides have been clearly observed in static structures, the rationale why arrestin engages with the entire receptor remains largely unclear. To investigate whether phosphopeptide binding to arrestin2 drives arrestin-GPCR complex formation, we analyzed the interactions of the isolated V2Rpp peptide, arrestin2, and β_1_AR in the presence and absence of orthosteric ligands by size exclusion chromatography (SEC, Figures 1A-D). The used β_1_AR construct is stabilized by point mutations and a truncation of the C-terminus after residue L367, but fully competent in agonist-induced G protein activation (19). Surprisingly, the V2Rpp-bound arrestin2 forms a stable complex at low micromolar affinity with carvedilol-bound β_1_AR even in the absence of chemical ligation between the V2Rpp peptide and the receptor. Carvedilol is a neutral antagonist (20) with almost absent G protein signaling, while it acts as an agonist for arrestin signaling (Table S1) (18), making it an ideal model for studying arrestin-GPCR interactions.

**Figure 1:**
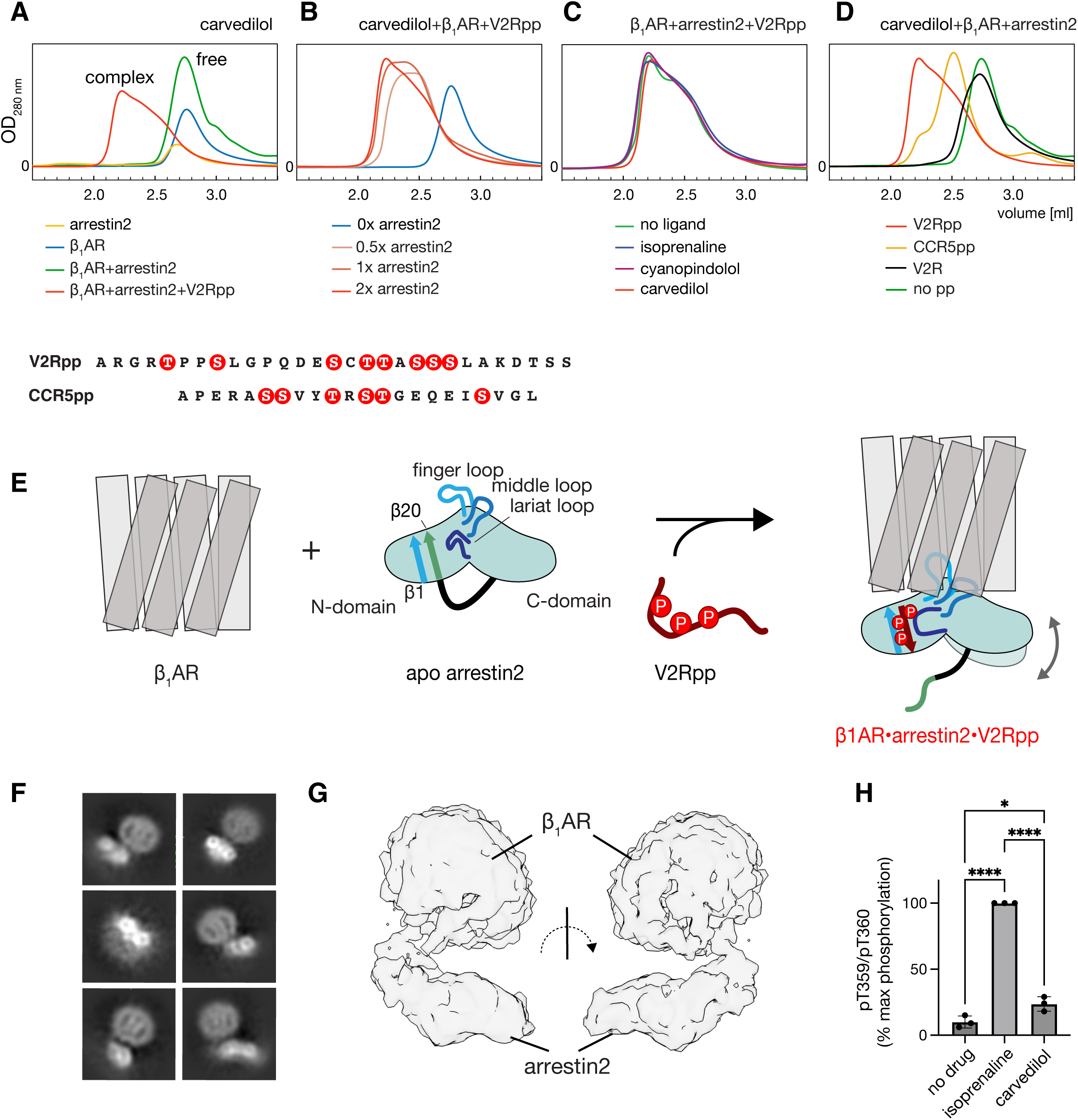
β_1_AR•arrestin2•V2Rpp and β_1_AR•arrestin2•CCR5pp complex formation. (A-D) SEC profiles of various mixtures of β_1_AR, orthosteric ligands, arrestin2 and phosphopeptides. In the individual panels, the constant part of the mixture is indicated at the top, whereas the varying part of the mixtures with color code is indicated at the bottom. The concentrations of individual components are given in Material and Methods. The V2Rpp and CCR5pp sequences are shown at the bottom with phosphosites marked by red circles. (E) Model of chemical equilibria leading to receptor-arrestin-phosphopeptide complex formation. (F) CryoEM 2D classes of carvedilol•β_1_AR•arrestin2•V2Rpp complex. (G) CryoEM 3D representation of carvedilol•β_1_AR•arrestin2•V2Rpp complex. (H) β_1_AR-V2Rpp phosphorylation detected in HEK293A cells by pT359/pT360 phospho-antibodies in the absence and presence of isoprenaline and carvedilol ligands. Individual phosphorylation amounts relative to isoprenaline, as well as their mean and standard deviation of the mean are shown for N=3 biological replicates; ****: P < 0.0001; *: P < 0.05.

The carvedilol•β_1_AR•arrestin2•V2Rpp complex exhibits a broad SEC elution curve with a maximum at 2.2 mL and a shoulder at 2.5 mL, which is shifted relative to the elution maximum of free β_1_AR at 2.7 mL. An SDS PAGE analysis of the SEC fractions showed a pure β_1_AR•arrestin2 complex with an approximate equimolar (1.0 ± 0.4) β_1_AR:arrestin ratio throughout the entire curve (Figure S1). The broad elution profile is very likely due to a dynamic equilibrium between free and bound molecules during gel filtration and a conformational exchange within the complex. As V2Rpp is not chemically ligated to the receptor, the interactions may not be very tight and also allow different arrestin orientations relative to the receptor.

Remarkably, the β_1_AR•arrestin2 complex formation does not depend on the presence of orthosteric ligands or their pharmacological properties, since V2Rpp-bound arrestin2 binds to apo β_1_AR as well as to the β_1_AR•isoprenaline (agonist) and β_1_AR•cyanopindolol (antagonist) complexes in the same way as to the β_1_AR•carvedilol complex (Figure 1C). Complex formation, albeit less efficiently, was also observed in the presence of the sixfold phosphorylated CCR5pp peptide derived from the C-terminus of the human chemokine receptor 5 (Figure 1D). The elution maximum of the CCR5pp complex occurs at 2.5 mL and hence is less shifted than for the V2Rpp complex. This reduced shift presumably arises from the lower affinity of CCR5pp (K_D_ = 45 μM) towards arrestin2 as compared to that of V2Rpp (K_D_ = 19 μM) (17) and from its lower phosphorylation (Figure 1). The weaker affinity may lead to a dynamic equilibrium during the flow on the SEC column that is closer to the free species. The absolute requirement of a phosphorylated peptide for the β_1_AR•arrestin2 complex formation is directly evident from experiments (Figure 1D) without a phosphopeptide or with the non-phosphorylated V2R peptide (V2R). In both cases, no complex is formed.

These results show that the arrestin2 recruitment to β_1_AR requires only the arrestin2 conformational change induced by the phosphopeptide binding and not the attraction of the positively charged arrestin2 to negatively charged phosphosites attached to the receptor. Furthermore, the recruitment is clearly independent of the specific β_1_AR functional state (Figure 2E).

**Figure 2:**
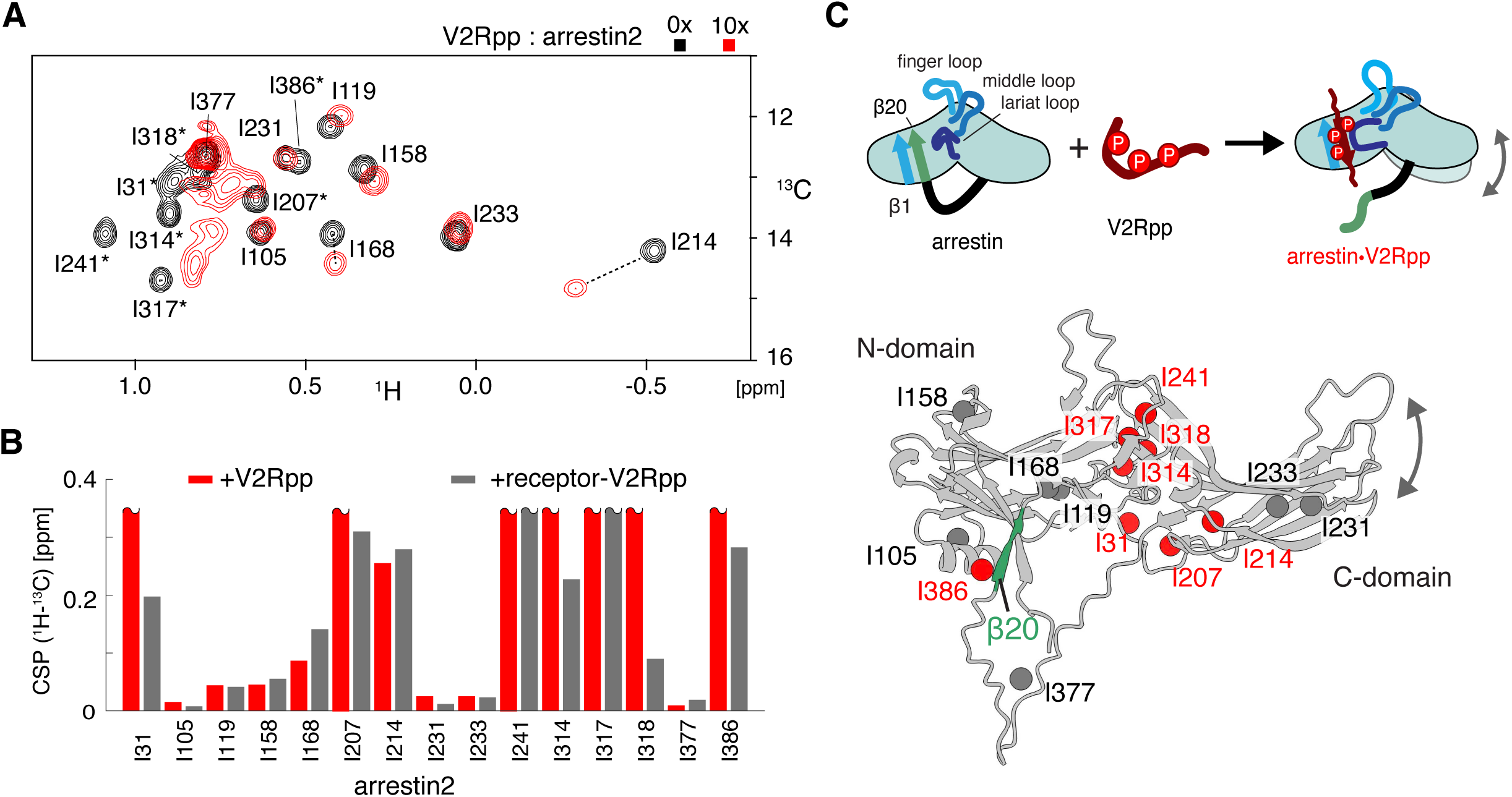
Arrestin2 activation by V2Rpp. (A) ^1^H-^13^C HMQC spectra of apo Ile-δ1-^13^CH_3_, ^2^H-labeled arrestin2 (black) and upon addition of a 10-molar equivalent of the V2Rpp (red). Resonances marked with an asterisk could not be assigned in the bound state. (B) Chemical shift perturbation (CSP) of arrestin2 isoleucine ^1^H_3_-^13^Cδ1 resonances upon addition of the V2Rpp phosphopeptide (red) or a β_2_AR-V2Rpp chimera (gray, data replotted from ref. (23)). CSP bars capped at 0.35 ppm correspond to resonances that could not be assigned or observed in the bound state due to severe broadening. (C) Schematics of arrestin activation: (top) The phosphopeptide displaces arrestin strand β20 (green) from the arrestin core and forms an intermolecular antiparallel β-sheet with strand β1 (blue). This induces changes in the arrestin loop conformations and the relative N- and C-domain orientation. (bottom) AlphaFold2 model of arrestin2 showing the N- and C-domains (gray). Isoleucine residues are indicated as spheres. Isoleucine residues colored in red undergo significant CSPs upon phosphopeptide binding.

The carvedilol•β_1_AR•arrestin2•V2Rpp complex proved sufficiently robust for low-resolution observation by cryoEM even in the absence of stabilizing antibodies (Figure 1F). The 2D classes reveal a well-formed complex with density visible both for β_1_AR and arrestin2 (Figures 1F and S2). While the relative flexibility of arrestin2 and β_1_AR prevented determination of a high-resolution structure, the data quality was sufficient to obtain a low-resolution 3D density map (Figure 1G, Table S2). Due to its limited resolution, the exact coordination between arrestin2 and β_1_AR remains unclear. A comparison to previously solved structures (Figure S3) shows that the density would allow the placement of a receptor core-engaged β_1_AR•arrestin2 complex, whereas placement of the receptor tail-engaged GCGR•arrestin2 complex leads to larger discrepancies.

As arrestin activation by interaction with a phosphorylated peptide appears as the main driving force of the receptor-arrestin interaction, we tested whether carvedilol has the ability to stimulate β_1_AR phosphorylation in HEK293A cells expressing β_1_AR fused to V2Rpp (Figures 1H and S4). Indeed, carvedilol increased β_1_AR-V2Rpp phosphorylation twofold relative to the basal signal in the absence of orthosteric ligands, which amounts to 20% of the phosphorylation induced by the full agonist isoprenaline. This agrees with the observed increased phosphorylation levels induced by carvedilol for a β_2_AR-V2Rpp fusion (18) as well as for β_1_AR with a native C-terminus (21) (Table S1).

### Activation by V2Rpp primes arrestin2 for complex formation with β_1_AR

To better understand the structural changes of arrestin2 induced by phosphopeptide binding in solution and in the absence of stabilizing antibody (Fab) fragments (8–14), we monitored the changes in the ^1^H-^13^C HMQC NMR spectrum of Ile-δ1-^13^CH_3_-labeled arrestin2 upon the addition of V2Rpp (Figure 2A). Ten-molar equivalents of the V2Rpp (corresponding to 95% of arrestin2 activation (17)) caused significant chemical shift perturbations of isoleucine residues 31, 207, 214, 241, 314, 317, 318, and 386 (Figures 2B and 2C). The change in Ile386 is a direct consequence of the phosphopeptide binding and displacement of the β20 strand. The other affected residues are located at the interface between N- and C-domain of arrestin2, which in the apo state is stabilized by the interaction of the arrestin core loops with the β strands of the N- and C-domain. The latter chemical shift changes are consistent with the observation that phosphopeptide binding induces a change in the conformations of the core loops and relative domain orientation by ∼20–25° as observed e.g. in V2Rpp•arrestin2 (16) and CCR5 phosphopeptide•arrestin2 (17) complexes as well as the V2Rpp•arrestin3 (22) complex, indicative of the very similar activation mechanism of the two non-visual arrestin isoforms. Apparently, the phosphopeptide-induced release of strand β20 together with the change in relative domain orientation and loop conformation prime arrestin for interaction with the receptor core.

It is interesting to note that the Ile-δ1-^13^CH_3_ chemical shift perturbations in arrestin2 induced by the addition of V2Rpp are highly similar to those reported for arrestin2 in the presence of a β_2_AR-V2Rpp chimera (Figure 2B) (23). This indicates that the binding of arrestin2 to the receptor does not change the phosphopeptide-induced active arrestin2 conformation much further. Our solution state NMR findings are supported by the almost identical structures of arrestin2 in phosphopeptide-bound (16) (PDB: 4JQI) and receptor-bound (8) (PDB:6TKO) forms.

### Arrestin recognizes both G protein binding-competent and -incompetent receptor states

To obtain insights into the β_1_AR conformation within the carvedilol•β_1_AR•arrestin2•V2Rpp complex under solution conditions, we carried out an extensive NMR analysis. Figure 3A shows the β_1_AR valine and tyrosine ^1^H-^15^N resonances of binary β_1_AR complexes with the antagonist carvedilol and the agonist isoprenaline, as well as the ternary complex of isoprenaline•β_1_AR with Nb80, which mimics G protein binding. When arrestin2•V2Rpp is added to carvedilol•β_1_AR, the resonances of the resulting quaternary β_1_AR•carvedilol•arrestin2•V2Rpp complex (Figure 3B, red) are broader and less intense, in agreement with its increased molecular weight and dynamic nature. Nevertheless, 23 ^1^H-^15^N β_1_AR resonances distributed throughout the receptor could be unambiguously detected in this complex. Their chemical shifts are very close to those of the binary β_1_AR•carvedilol complex, but strongly differ from the isoprenaline•β_1_AR•Nb80 complex (Figure 3C,D). Thus, the entire receptor within the quaternary arrestin2 complex clearly adopts the same G protein-inactive conformation as in the β_1_AR•carvedilol complex.

**Figure 3:**
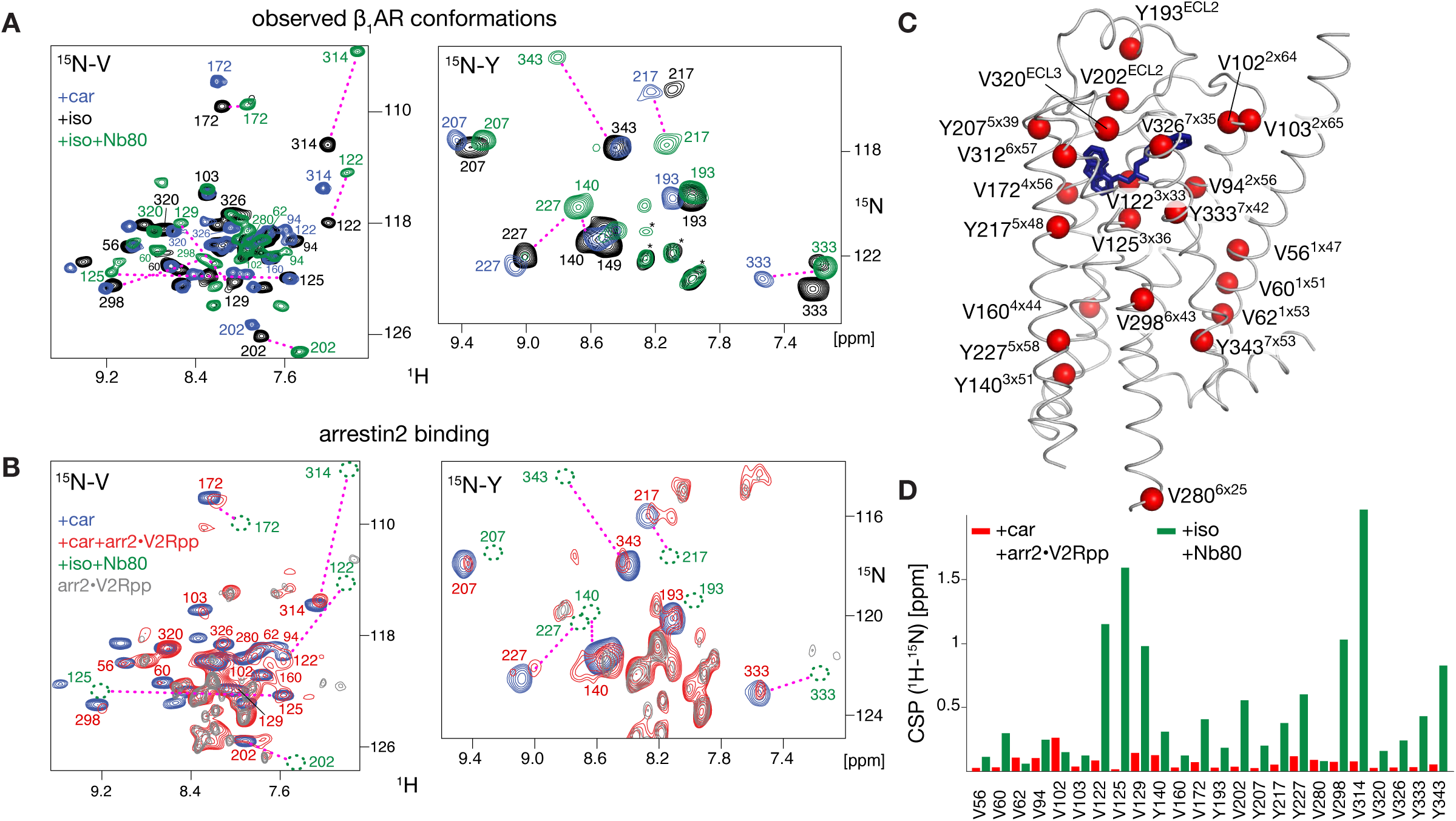
Determination of the β_1_AR functional state by ^1^H-^15^N valine and tyrosine NMR. (A) Superposition of ^1^H-^15^N TROSY spectra of ^15^N-valine-(left) and ^15^N-tyrosine-labeled β_1_AR (right) in binary complex with carvedilol or isoprenaline and ternary complex with isoprenaline and Nb80. Resonances are marked with assignment information. Resonances connected by a dashed magenta line represent residues with two clearly distinguishable resonances for the inactive (carvedilol•β_1_AR complex) and active (isoprenaline•β_1_AR•Nb80 complex) conformation. (B) as (A) Spectra of the quaternary carvedilol•β_1_AR•arrestin2•V2Rpp complex superimposed onto spectra of carvedilol•β_1_AR and arrestin2•V2Rpp (^15^N natural abundance). The resonance positions of the G protein-active isoprenaline•β_1_AR•Nb80 complex of panel (A) are shown in comparison by green dashed ellipses. Resonances not assigned to any β_1_AR residue in (A) and (B) are marked by asterisks. Notably, the ^15^N-natural-abundance resonances of arrestin2 become visible in the spectra of the carvedilol•β_1_AR•arrestin2•V2Rpp complex. (C) Structure of isoprenaline•β_1_AR complex (PDB 2Y03) with all valine and tyrosine residues identified by NMR to be in the inactive state for the carvedilol•β_1_AR•arrestin2•V2Rpp complex (D). Chemical shift perturbation (CSP) of ^1^H-^15^N resonances of carvedilol•β_1_AR•arrestin2•V2Rpp (red) or isoprenaline•β_1_AR•Nb80 (green) complexes relative to the carvedilol•β_1_AR complex.

We also tested a further β_1_AR construct (β_1_AR^V129I^) where the thermostabilizing V129 is back-mutated to the native I129. In comparison to β_1_AR, the conformational equilibrium of the binary isoprenaline•β_1_AR^V129I^ complex is strongly shifted from the G protein-preactive to the G protein-active conformation, while β_1_AR^V129I^ complexes with antagonists remain unchanged (24, 25). However, also for the isoprenaline•β_1_AR^V129I^•arrestin2•V2Rpp complex no resonances corresponding to the G protein-active conformation could be detected in the ^1^H-^15^N tyrosine spectrum (Figure S5). Instead, the tyrosine resonances appear shifted towards the G protein-preactive conformation.

Notably, the tyrosine ^1^H-^15^N resonances monitor the conformation of the two conserved tyrosines Y^5×58^ and Y^7×53^ (superscripts indicate Ballesteros–Weinstein numbering (26)), which are critical for receptor activation (27) (Figure 4). In the active G protein-bound state, the sidechains of both residues form a water-mediated bridge (YY-bridge closed, Figure 4A) as observed e.g. in adrenergic receptors (β_1_AR – PDB 6H7J (28) and 6IBL (8), β_2_AR – PDB 7BZ2 (29)). This YY-bridge stabilizes the swung-out positions of TM5 and TM6, which allow the accommodation of the G protein. In the inactive state, the YY-bridge is open and the opening of the intracellular helical bundle is reduced (Figure 4A). The Y^7×53^ side chain is then rotated and interacts with the side chain amide of N^7×49^ within the NPxxY motif, whereas the Y^5×58^ side chain faces towards the intracellular opening of the transmembrane helix bundle.

**Figure 4:**
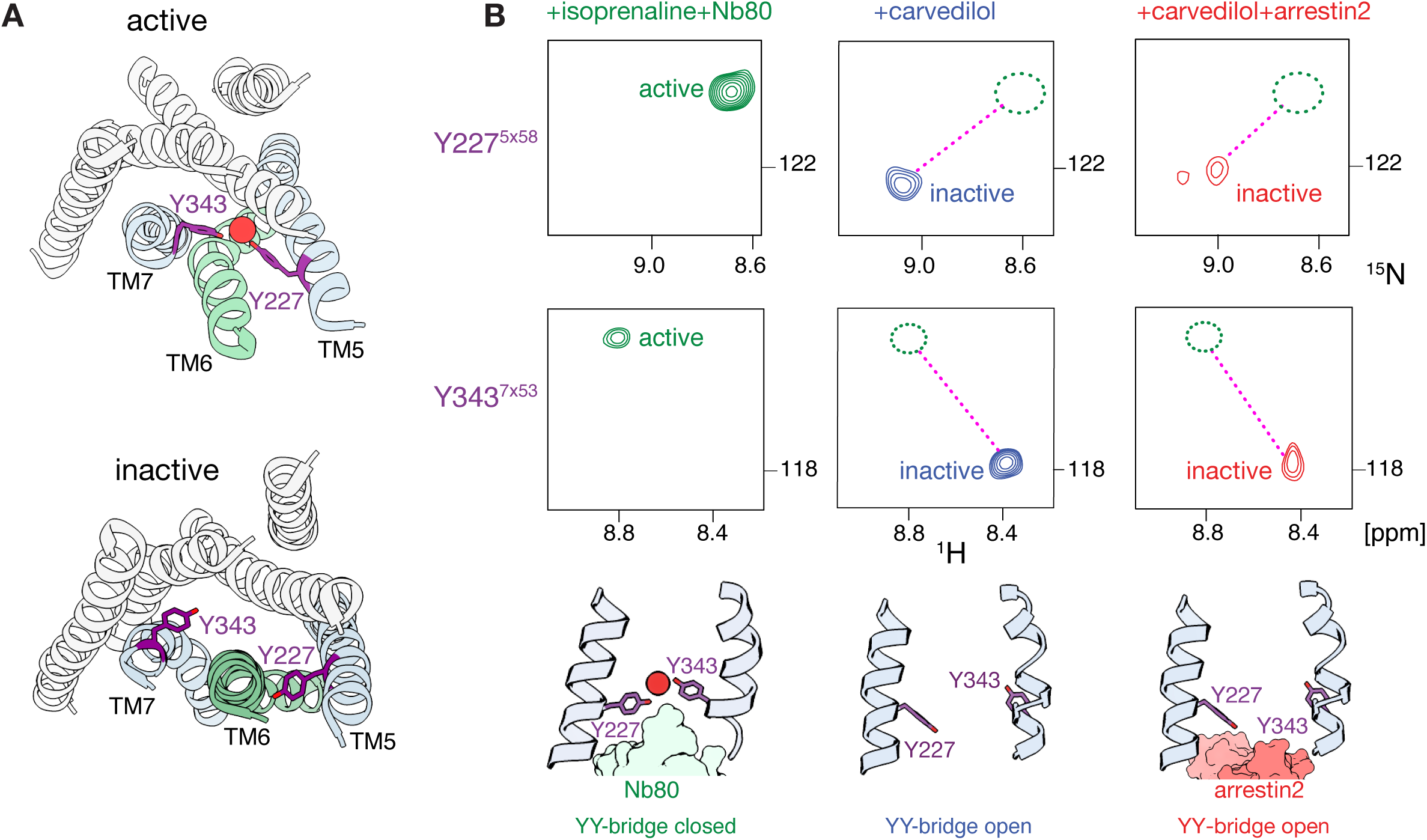
Arrestin2 binds to β_1_AR in G protein-inactive conformation. (A) G protein-active and -inactive conformations of Y227^5×58^ and Y343^7×53^ in β_1_AR. Top: active conformation of TM5 and TM7 (light blue) with water-mediated (red sphere) bridge between Y227 and Y343 (YY-bridge closed, isoprenaline•β_2_AR•Nb80 complex, PDB 6H7J). Bottom: inactive TM5/TM7 conformation with Y227 and Y343 separated (YY-bridge open, carvedilol•β_2_AR complex, PDB 6PS3). The carvedilol•β_2_AR structure was used to depict the open YY-bridge because Y227 is mutated to alanine in the corresponding β_1_AR structure. (B) Selected regions of the ^1^H-^15^N TROSY spectra showing the β_1_AR Y227 and Y343 resonances in different complexes and functional states. The green dashed ellipses indicate resonances absent in the active conformation. The structures at the bottom show the β_1_AR YY-bridge conformations observed by NMR [β_1_AR+isoprenaline+Nb80: closed (PDB 6H7J), β_1_AR+carvedilol: open (carvedilol•β_2_AR complex, PDB 6PS3), β_1_AR+carvedilol+V2Rpp+arrestin2: open (modeled using the receptor from the carvedilol•β_2_AR complex, PDB 6PS3, and arrestin2 from the formoterol•β_1_AR-V2Rpp•arrestin2 complex, PDB 6TKO)].

For the binary β_1_AR complexes with the antagonist carvedilol or the agonist isoprenaline, the ^1^H-^15^N resonances of Y227^5×58^ and Y343^7×53^ (Figure 3A and 4B) are very similar and indicate an open YY-bridge. In contrast, both tyrosines undergo strong chemical shift changes in the active ternary complex with isoprenaline and Nb80, indicative of the closed YY-bridge. In the β_1_AR•carvedilol•arrestin2•V2Rpp complex the resonances are weaker, but clearly are in the position of the open YY-bridge. Hence the NMR data indicate that arrestin2 recognizes β_1_AR in an inactive conformation with an open YY-bridge that would exclude G protein binding. The inability to bind to the G protein is confirmed by the NMR spectra, which showed no changes and indication of complex formation for carvedilol-bound β_1_AR when either the G protein mimicking Nb80 (Figure S6A) or miniGs (Figure S6B) were added.

In addition to defining the G protein activation state of the receptor, the observed ^1^H-^15^N resonances offer deeper insights into specific conformations related to the arrestin bias of carvedilol. Arrestin signaling has been associated in particular with TM7 by DEER (30) and ^19^F-NMR (31) observations as well as molecular dynamics simulations (32). Figure S7 shows a comparison of ^1^H-^15^N tyrosine and valine resonances of β_1_AR in complex with the very weak G protein agonist cyanopindolol (Table S1) and carvedilol. Not surprisingly, the strongest chemical shift changes are observed around the orthosteric binding site for residues V125^3×36^, V309^5×54^, and Y333^7×42^. Interestingly, the resonances of the YY-bridge residues Y227^5×58^ and Y343^7×53^ show no particular variations and are in the open YY-lock position. However, several residues in the cyanopindolol•β_1_AR complex give evidence of a dynamical equilibrium: (i) Y333^7×42^shows a splitting into two resonances: one that is close to the resonance of the G protein-active conformation of the isoprenaline•β_1_AR•Nb80 complex and another that coincides with the single resonance of the inactive carvedilol•β_1_AR complex. (ii) Y140^3×51^, located close to the intracellular side of TM3, has a resolved resonance at the same location as in isoprenaline•β_1_AR•Nb80 complex (Figure 3A). A second Y140^3×51^ resonance may also be present in the position of the inactive carvedilol•β_1_AR complex, but is obscured by overlap with Y149. (iii) the V122^3×33^ resonance is missing in the cyanopindolol•β_1_AR complex, apparently due exchange broadening. In contrast, all these residues have well resolved single resonances in the carvedilol•β_1_AR complex. Thus, carvedilol tightly locks the receptor in a single inactive conformation. This can be rationalized by the filling of a receptor pocket located between TM3 and TM7 by the bulky anisole tail moiety of carvedilol, which is left empty by the shorter cyanopindolol (Figure S7D). The stabilization of the inactive conformation by the increased packing of this part of the orthosteric pocket may contribute to carvedilol’s arrestin-biased agonism.

### Y227 and Y343 are not required for arrestin2 binding

The conserved Y227^5×58^ and Y343^7×53^ tyrosines are essential for G-protein binding (19, 27). To investigate their significance for the arrestin interaction, we generated a β_1_AR mutant (β_1_AR^Y227A,Y343L^) lacking these residues. Remarkably, the β_1_AR^Y227A,Y343L^ mutant also forms a stable complex with the V2Rpp-activated arrestin2 as observed by SEC experiments (Figure 5A). The sharper SEC profile and slightly reduced elution volume of this β_1_AR^Y227A,Y343L^•arrestin2•V2Rpp complex relative to the β_1_AR•arrestin2•V2Rpp complex indicate an even higher stability. As expected, no complex formation of β_1_AR^Y227A,Y343L^ was observed in the absence of V2Rpp (Figure 5A).

**Figure 5:**
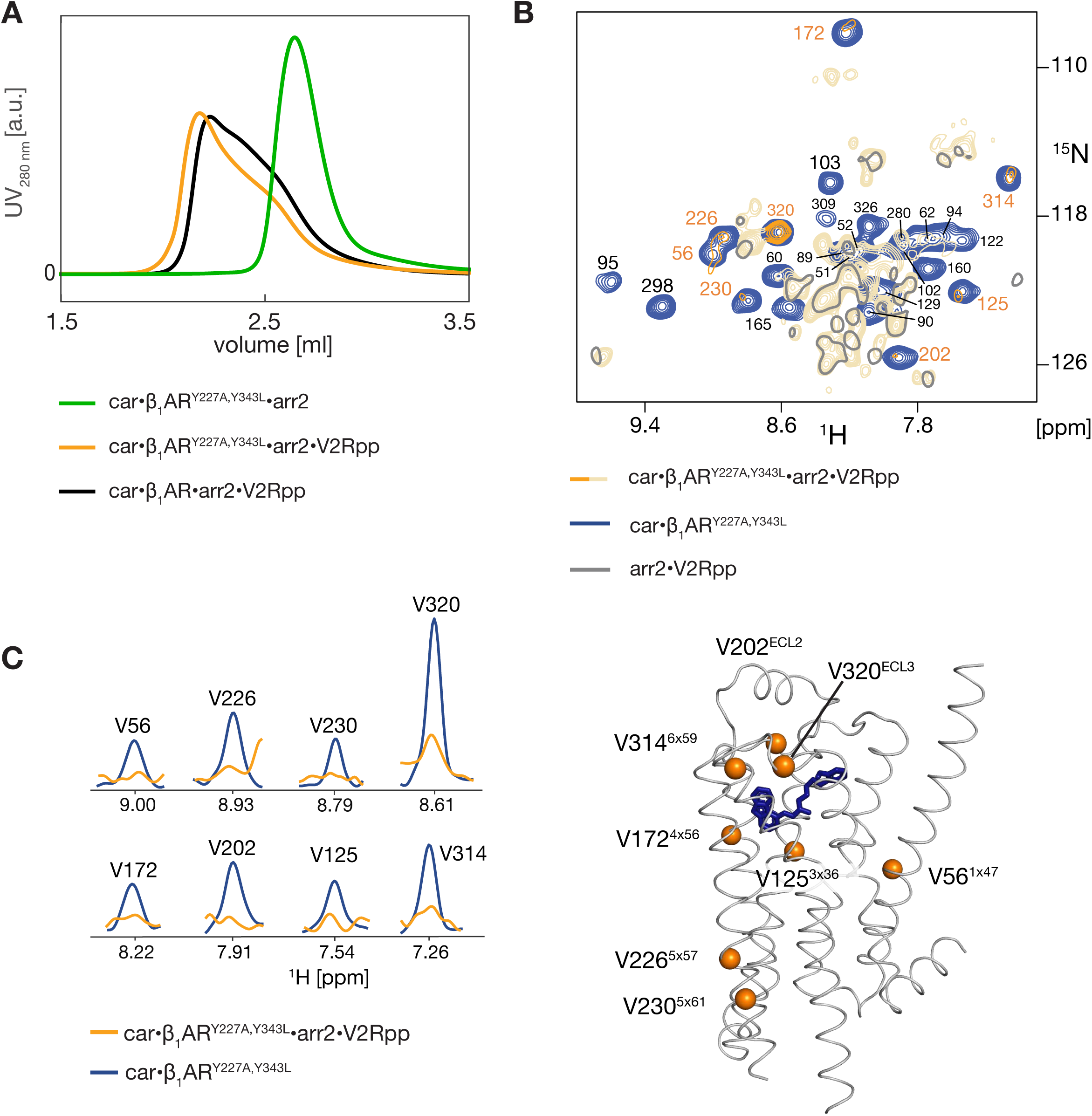
The G protein binding-incompetent mutant β_1_AR^Y227A,Y343^ binds arrestin2. (A) SEC profiles of mixtures of carvedilol+β_1_AR^Y227A,Y343L^+arrestin2 (green), carvedilol+β_1_AR^Y227A,Y343L^+arrestin2+V2Rpp (orange), and carvedilol+β_1_AR+arrestin2+V2Rpp (black). The concentrations of individual components are given in Methods. (B) Superposition of ^1^H-^15^N TROSY spectra of ^15^N-valine-labeled carvedilol•β_1_AR in absence (blue) and presence of a 1.5-molar equivalent arrestin2•V2Rpp (orange). Valine resonances are marked with assignment information. The severe broadening of the β_1_AR valine resonances upon addition of arrestin2•V2Rpp indicates complex formation. Due to the line broadening, the sensitivity of the ^15^N-valine detection is low in the quaternary complex, and natural abundance ^15^N arrestin2 resonances (light orange) become dominant. The ^15^N natural abundance TROSY spectrum of an arrestin2:V2Rpp 1:5 molar mixture (grey, single contour lines) is shown as a control. (C) 1D traces through selected ^1^H-^15^N valine resonances of carvedilol•β_1_AR (blue) carvedilol•β_1_AR•arrestin2•V2Rpp (orange) spectra in panel (B) indicating the line broadening due to complex formation. (D) Valine residues identified by NMR in the inactive state of the carvedilol•β_1_AR•arrestin2•V2Rpp complex shown as orange spheres within the structure of carvedilol•β_1_AR (PDB 4AMJ).

The conformational state of β_1_AR^Y227A,Y343L^ in complex with arrestin2 was also characterized by NMR (Figures 5B-C). For the binary complex of ^15^N-valine labeled β_1_AR^Y227A,Y343L^ with carvedilol all expected 28 ^1^H-^15^N valine resonances are detected with high sensitivity in a ^1^H-^15^N TROSY spectrum (Figure 5B). Upon addition of V2Rpp and arrestin2, all the valine resonances are dramatically broadened (Figure 5C), as a result of the complex formation. The strongest ^1^H-^15^N resonances in this spectrum result from the natural ^15^N abundance of V2Rpp and arrestin2. However, all detected β_1_AR^Y227A,Y343L^ valine resonances are in positions corresponding to the inactive carvedilol binary-complex. This is expected since β_1_AR^Y227A,Y343L^ is unable to reach the active conformation (19).

## Discussion

### Phosphopeptide interactions and GPCR phosphorylation

The currently available structures show arrestin2 in core-engaged complexes only with phosphorylated GPCRs (8, 14, 10, 13, 9, 33) that are in a G protein-active conformation. However, MD simulations (32) suggest that arrestin may also form core-engaged complexes with receptor conformations that are inaccessible for the G protein. Thus, the precise requirements of arrestin recruitment to GPCRs have remained unclear. To clarify these requirements, we have investigated the molecular details of arrestin2 interaction with the β_1_AR. Strikingly, stable complexes of β_1_AR and V2Rpp-activated arrestin2 could be generated in the absence of chemical ligation of V2Rpp with the receptor and without stabilizing antibody fragments. The interaction with the receptor itself does not change the conformation of the activated arrestin2 appreciably. The independence of the interaction from the phosphopeptide-receptor ligation implies that the β_1_AR•arrestin2 complex formation is not dominated by the electrostatic attraction of arrestin2 to the phosphorylated receptor.

We have previously determined the affinity of phosphopeptides towards arrestin2 (17). The affinity depends only moderately on the phosphosite location, but strongly increases with their number, with K_D_s larger than millimolar observed for a non-phosphorylated CCR5 C-terminal peptide, ∼400 μM for a 3-fold phosphorylated CCR5 C-terminal peptide, and ∼20 μM for the 8-fold phosphorylated V2Rpp, respectively. In agreement with this trend, the strength of arrestin recruitment and interactions is weaker for so-called arrestin-class A GPCRs, which have lower phosphosite density than arrestin-class B receptors (17, 34). However, membrane-associated phosphoinositides (PIPs) can enhance arrestin recruitment for class A receptors by charge interactions with the positive arrestin C-lobe (35).

In contrast to the arrestin-phosphopeptide affinity, arrestin activation as assayed by Fab30 binding depends on the exact positions of the phosphosites with a pXpp motif playing a central role (17). This pXpp motif is specifically recognized by arginine and lysine residues within the arrestin N-domain. The arrestin activation leads to the release of strand β20 and changes in the core loop conformations, which enable the subsequent interaction with the receptor core. Thus, the arrestin activation and strong receptor core interaction are apparently induced by specific receptor phosphorylation patterns, whereas the overall recruitment of arrestin to the receptor within the membrane is modulated by electrostatic interactions between the phosphosites or other receptor-associated negative charges and the largely positive arrestin.

As the presence of a phosphorylated peptide, albeit not necessarily covalently attached, is a prerequisite for arrestin2 binding to β_1_AR, we verified that the arrestin-biased ligand carvedilol is able to induce the phosphorylation of a β_1_AR-V2Rpp fusion in HEK293A cells. This agrees with cellular assays on a β_2_AR-V2Rpp fusion (18) or wild-type β_1_AR (21), in which phosphorylation by GRK and subsequent arrestin recruitment was induced by agonists and carvedilol (Table S1). Corroborating our observations for β_1_AR^Y227A,Y343L^, also a G protein-inactive β_2_AR-V2Rpp mutant (T68^2×39^F, Y132^3×51^G, Y219^5×58^A) induced the same arrestin signaling and phosphorylation profile as the wild-type construct (18).

### Independence from orthosteric ligand

The complex formation between β_1_AR and V2Rpp-activated arrestin2 is independent of receptor stimulation by an orthosteric ligand and its G protein agonistic or antagonistic properties. This agrees with previous findings that an arrestin core mutation (R169E in arrestin2), which activates arrestin in the absence of phosphopeptides, promotes interaction with GPCRs (36) and that such a constitutively active arrestin induces agonist-independent internalization of the 5-HT_2A_ serotonin receptor (37).

### The carvedilol•β_1_AR•arrestin2•V2Rpp complex is in a G protein-inactive conformation

To determine the conformation of β_1_AR under solution conditions in the arrestin2-bound state, we compared 23 distinct ^1^H-^15^N NMR resonances of tyrosine and valine residues spanning the entire receptor in the carvedilol•β_1_AR•arrestin2•V2Rpp complex, the carvedilol•β_1_AR complex and the isoprenaline•β_1_AR•Nb80 complex, as a mimic for the G protein-bound state. The chemical shifts clearly show that the arrestin2-bound carvedilol•β_1_AR is in an identical conformation as the inactive binary carvedilol•β_1_AR complex and not in the G protein-bound active state. In particular, the conserved YY-bridge is open, whereas a closed YY-bridge is observed in the Nb80•β_1_AR, the G protein•β_2_AR, as well as the β_1_AR•arrestin2 complex structures. Supporting this observation, mutations of the Y227^5×58^ and Y343^7×53^ residues, which are essential for G protein binding, do not impair V2Rpp-activated arrestin2 recruitment by the carvedilol-bound β_1_AR (Figure 4).

### Arrestin promiscuity

The G protein-inactive receptor conformation in the carvedilol•β_1_AR•arrestin2•V2Rpp complex contrasts with the G protein-active conformation with swung-out TM5 and TM6 in all currently solved structures of core-engaged GPCR-arrestin complexes (β_1_AR (8), M2R (14), CB1R (9), NT1R (7), V2R (10), 5HT2bR (13), rhodopsin (6)). This proves that arrestin is able to bind to both G protein-active and -inactive receptor conformations and explains the observed independence of the arrestin2 binding from the orthosteric ligand in the SEC assays (Figure 1). The promiscuity of arrestin towards GPCRs in different functional states is also in agreement with MD molecular dynamics simulations of the AT_1_R receptor (32), where two major signaling conformations were observed, one of which coupled to both arrestin and a G protein, whereas the other almost exclusively to arrestin. The switch between both conformations involved a complete reorientation of the Y^7×53^ sidechain.

### Conclusion

The independence of arrestin binding from the specific receptor conformation provides not only an explanation why only two non-visual arrestin isoforms are capable of regulating the hundreds of non-visual GPCRs (3). It may also explain the distinct pharmacological profile of the beta-blocker carvedilol as an arrestin-biased agonist. As the activated arrestin recognizes inactive and active receptors with similar efficiency, the arrestin-biased signaling of carvedilol appears to be caused by the conformational exclusion of G protein, rather than by a specific receptor conformation that selects arrestin. By tightly modulating the conformational state of β_1_AR, carvedilol primarily prevents G protein binding, while still promoting receptor phosphorylation, thereby allowing activated arrestin to engage effectively (Figure 6). These molecular insights into the mechanism of arrestin activation and recruitment to GPCRs may help to develop therapies that target GPCR signaling pathways in a selective manner.

**Figure 6:**
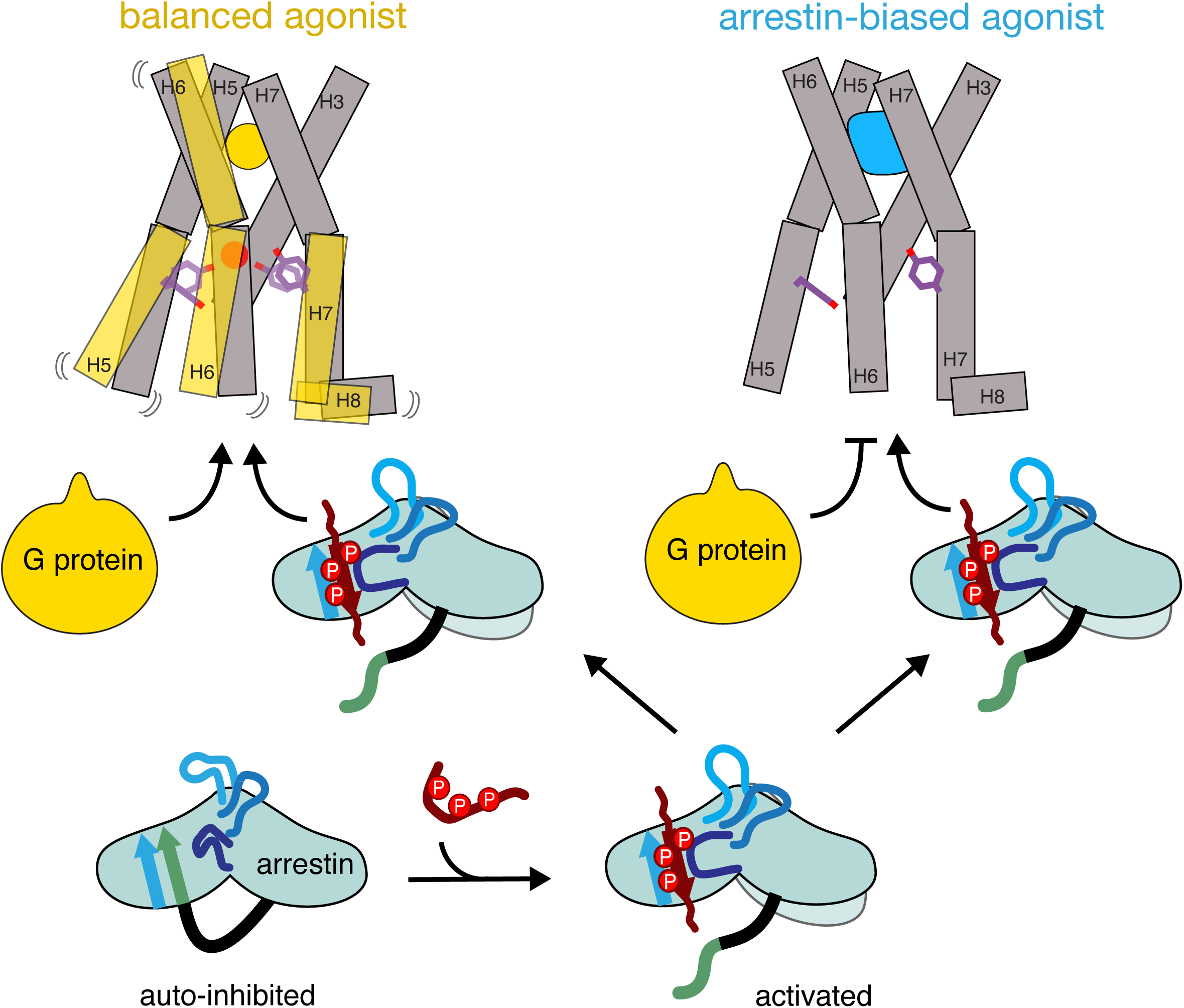
Structural basis of the carvedilol arrestin-biased agonism. Activation of auto-inhibited arrestin is induced by the addition of phosphopeptide that primes arrestin for receptor engagement. Left: Balanced agonists promote increased dynamics of the intracellular side of the receptor with open and closed conformations of the helical bundle and the YY-bridge. This allows both G protein and arrestin to bind to the receptor. Right: arrestin-biased agonists, such as carvedilol, stabilize a G protein-inactive state, with the intracellular opening closed and the YY-bridge open, to which only arrestin but not to G proteins can bind.

## Materials and methods

### Protein expression and purification

The expression of the thermostabilized unlabeled, ^15^N-valine-labeled, or ^15^N-tyrosine-labeled turkey β_1_AR G protein binding-competent construct (extinction coefficient ε_280_ = 68410 M^-1^cm^-1^), the more native β_1_AR^V129I^ construct, and the G protein binding-incompetent variant β_1_AR^Y227A,Y343L^ in baculovirus-infected Sf9 cells, purification, binding of ligands, and assignments were carried out as described previously (19, 27, 38).

Unlabeled arrestin2 (residues 1-393, ε_280_ = 19450 M^-1^cm^-1^) expression and purification were performed as described before (17). For the preparation of Ile-δ1-^13^CH_3_-labeled arrestin2, the *E. coli* (BL21) cells were grown in D_2_O M9 minimal medium supplemented with 80 mg per liter of 2-ketobutyric acid-4-^13^C,3,3-d_2_ sodium salt hydrate following previously reported procedures (39).

The plasmids for Nb80 and miniGs were generous gifts by Dr. Jan Steyaert and Dr. Chris Tate, respectively. Nb80 and miniG_S_ were produced according to described procedures (40, 41).

### Peptide synthesis

The phosphorylated V2R peptide (V2Rpp, ARGRpTPPpSLGPQDEpSCpTpTApSpSpSLAKDTSS) corresponding to the last 29 residues of the Vasopressin receptor 2 and the phosphorylated CCR5 peptide (CCR5pp, APERApSpSVYpTRpSpTGEQEISpVGL) corresponding to the last 22 residues of the CC chemokine receptor 5 were obtained from the Tufts University Core Facility for peptide synthesis.

### Size exclusion chromatography

SEC binding assays were performed on an UltiMate 3000 HPLC system using a self-packed 4.2-mL S200 10/300 SEC column (length 25 mm, diameter 4.6 mm) pre-equilibrated with SEC buffer (20 mM Tris-HCl, 100 mM NaCl, ∼40 mM decylmaltoside (DM), 0.02% NaN_3_, pH 7.5). Samples containing 20 μM β_1_AR (1x), 40 μM arrestin2 (2x), 100 μM V2Rpp (5x), 200 μM CCR5pp (10x) and 1 mM (carvedilol, cyanopindolol) or 2 mM (isoprenaline) ligand were prepared as 270 μL volumes in SEC buffer. The SEC buffer was supplemented with 10 μM carvedilol, 10 μM cyanopindolol or 1 mM isoprenaline, respectively. Control samples were prepared at the same concentrations as in the sample mixtures with the exception of the arrestin2 control (20 μM).

### Cryo-EM sample preparation and grid freezing

The carvedilol•β_1_AR•arrestin2•V2Rpp complex for cryo-EM analysis was prepared similarly to the samples for the SEC-binding assay. Samples were prepared as 270 μL volumes and contained 50 μM receptor, 100 μM arrestin2, 250 μM V2Rpp, 1 mM carvedilol and 0.1 % DM. The samples were incubated for 30 min on ice before injecting onto the 4.2 mL SEC column preequilibrated with cryo-EM buffer (20 mM HEPES, 100 mM NaCl, 0.01 % LMNG, pH 7.5, 10 μM carvedilol). The SEC fractions were checked using SDS-PAGE, correct fractions pooled and concentrated to 1.4 mg/ml.

### Cryo-EM data collection and structure determination

3-µl aliquots of the sample were deposited onto glow-discharged Quantifoil^TM^ R1.2/1.3 (Cu) or R2/1 (Gold) grids and rapidly frozen in liquid ethane using a Leica EM GP2 plunger adjusted to +4°C and 80% humidity, with a blotting time of 3-5 seconds. Data acquisition as Electron Event Recordings (EER) was performed on a TFS Titan Krios G4 microscope, equipped with a cold-FEG electron source and Falcon4i camera (DCI-Lausanne), detailed in Table S2.

Recorded EER files were processed in RELION 4.0 (42, 43). Motion correction was performed using RELION’s implementation of Motioncor2, with 32 EER fractions per movie frame. CTF estimation was conducted using CTFFIND 4.1.14 (44), with selection criteria based on a maximum CTF resolution of better than 6 Å and defocus between 0.5 to 2.5 µm. Particle picking utilized template matching or blob picking methods, yielding a large particle dataset (Table S2). The particle stacks were imported to cryoSPARC (45) and rounds of 2D classification were carried. Good 2D classes were used for ab initio model generation. The volumes were visualized in ChimeraX (46).

### NMR experiments

NMR samples were prepared with typical receptor concentrations of 100−200 μM in 20 mM Tris-HCl, 100 mM NaCl, ∼40 mM decylmaltoside (DM), 0.02% NaN_3_, 5% D_2_O, pH 7.5. Binary complexes with the various orthosteric ligands were formed by adding 2 mM ligand. For isoprenaline, 20 mM sodium ascorbate was added to prevent oxidation. Ternary complexes were formed by adding a 1.2-molar equivalent of Nb80, a 1.5-molar equivalent of miniG_S_, or a 1.5-molar equivalent of arrestin2 together with a 5-molar equivalent of V2Rpp, respectively.

^1^H-^15^N TROSY NMR experiments were recorded on Bruker AVANCE 900 MHz or AVANCE 600 MHz spectrometers equipped with TCI cryoprobes at 304 K using sample volumes of ∼250 μL in Shigemi microtubes. ^1^H-^15^N TROSY experiments were recorded as 80 (^15^N) × 1024 (^1^H) complex points with acquisition times of 16 ms (^15^N) and 42 ms (^1^H). For optimal sensitivity, the ^1^H-^15^N transfer time was reduced to 3.0 ms and a 1 s interscan delay was used. Typical total experimental times were 48 h.

A titration of 50 μM Ile-δ1-^13^CH_3_, ^2^H-labeled arrestin2 with V2Rpp up to a concentration of 500 μM was monitored by ^1^H-^13^C HMQC spectra. The assignments of the methyl groups were transferred from BMRB ID:51131 (23).

All NMR spectra were processed with NMRPipe (47) and evaluated with CCPNmr Analysis (48) or NMRFAM-SPARKY (49). 1D-NMR traces were generated using the nmrglue (50) module in the PyCharm environment (JetBrains, PyCharm 2020.1.2).

Chemical shift perturbations (CSP) were calculated using Equation 1

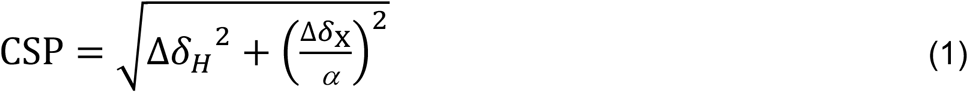

where Δδ_H_ and Δδ_X_ represent the chemical shift differences of the proton and the directly attached heteronucleus (^13^C or ^15^N), respectively. The scaling factor α was set to 5.7 or 5.0 for ^13^C or ^15^N, respectively. CSPs larger than the average CSP calculated from all resonances were considered significant.

### Structural analysis

Molecular visualization was carried out with UCSF ChimeraX (51).

### Cellular assays

HEK293A cells were grown in high-glucose Dulbecco’s modified Eagle’s medium (Sigma-Aldrich) with 10% fetal bovine serum (FBS, Biowest), 2 mM L-glutamine, 100 U ml^−1^ penicillin G and 100 ng ml^−1^ streptomycin, 1 mM sodium pyruvate at 37 °C and 5% CO_2_.

To assess the level of phosphorylation of the receptor, cells were plated into six-well plates to reach 70% confluency. Cells were transfected with 0.3 µg of each arrestin2-YFP and β_1_AR-V2Rpp plasmid DNA (GenScript) complexed with Helix-IN transfection reagent (OZ Biosciences). Transfection was carried out according to the manufacturer’s recommendation. Post 24 hours of transfection, cells were treated with 100 µM of isoprenaline (positive control) and 10 µM of carvedilol for 30 minutes. Cells were lysed using RIPA buffer (50mM Tris pH 8, 150 mM NaCl, 5 mM EDTA, 0.5% sodium deoxycholate, 0.1 % SDS, 1 % Triton X-100) supplemented with 1x Halt’s protease and phosphatase inhibitor cocktail.

Phosphorylation and total amount of flag-tagged β_1_AR-V2Rpp were quantified by western blotting using chemiluminescence. For this, 20 μL of the respective fractions were diluted with 5 μL SDS–PAGE buffer (125 mM Tris–HCl pH 6.8, 4% (w/v) SDS, 20% (v/v) glycerol and 0.01% (w/v) bromophenol blue) and loaded onto Mini-PROTEAN TGX™ 4–20% precast gels (Bio-Rad). As a reference, 3 μL of Precision Plus Protein™ dual color standard weight ladder (#1610374, Bio-Rad) were also loaded onto the gel, and the proteins separated for 30 min at 200 V. Proteins were transferred to 0.2 μm nitrocellulose membranes using the Trans-Blot Turbo™ transfer system (Bio-Rad). Western-blot membranes were blocked with 1% BSA in Tris-Buffered Saline-Tween (TBST) for 1 h and then incubated with monoclonal HRP anti-flag tag antibody (1:5000 dilution, Thermo Fisher) or with primary (rabbit polyclonal) phospho-V2R [pT359/pT360] antibody (1:2000 dilution, 7TM antibodies) followed by HRP-coupled anti-rabbit secondary antibody (1:5000 dilution, Thermo Fisher) for 1 hour each. Blots were then washed three times for 5 min with TBST and developed using the Lumi-Light Western Blotting Substrate (Hoffmann-La Roche). The chemiluminescence signal was detected using a Fusion FX6 reader (Vilber) at 600 dpi resolution. Individually selected bands of the obtained images from the western blot membranes were quantified by ImageJ (52) after background subtraction using the rolling ball radius method (90 pixels). Phosphorylation levels were normalized using the respective total amount of expressed flag-tagged β_1_AR-V2Rpp.

### Statistics

A statistical analysis of cellular receptor phosphorylation was performed using GraphPad Prism 10.1.1 (GraphPad Software, Inc., San Diego, CA, USA). The normality of the individual ligand-induced phosphorylation results was tested using the Shapiro-Wilk normality test and the data were cross-compared using an ordinary one-way ANOVA test.

## Data availability

The Cryo-EM density map for β_1_AR in complex with carvedilol, arrestin2, and V2Rpp was deposited to the Electron Microscopy Data Bank (EMDB) with accession code EMDB-50623 and the raw cryo-EM images were deposited to the Electron Microscopy Public Image Archive (EMPIAR) with accession code EMPIAR-12139. NMR spectra and peak assignments are accessible through the Biological Magnetic Resonance Bank (BMRB) with accession codes XXX the submission is in progress XXX.

## Acknowledgments and funding sources

This work was supported by the Swiss National Science Foundation (grants CRSK-3_195592 to L.A.A., and 31-201270 and IZLIZ3-200298 to S.G, 310030-188548 to HS, and 310030-197779 to A.S), and by a Fellowship for Excellence by the Biozentrum Basel International PhD Program to I.P. We gratefully acknowledge Drs. Timothy Sharpe and Tobias Mühlethaler (Biozentrum Biophysics Facility) for expert help with the biophysical characterization of β_1_AR, Drs. Mohamed Chami, Carlos Fernandez Rodriguez, and Carola Alampi (Biozentrum BioEM lab) for cryo-EM initial screenings, the Dubochet Center for Imaging Lausanne (a joint initiative from EPFL, UNIGE, UNIL, UNIBE) with the assistance of Drs. A. Myasnikov, B. Beckert, S. Nazarov, I. Mohammed, and E. Uchikawa for cryo-EM data collection, as well as Dr. Timm Maier for helpful discussions.

## Author contributions

L.A.A., I.P., and S.G. conceived the study. L.A.A. expressed and purified β_1_AR and β_1_AR^Y227A,Y343L^, and recorded and analyzed NMR data on those proteins. I.P. expressed and purified arrestin2 and recorded and analyzed NMR data of arrestin2 with V2Rpp. A.G. recorded ^15^N-valine-β_1_AR•carvedilol spectra. I.P. and L.A.A. designed SEC experiments and prepared complexes for EM. M.T. prepared EM grids and collected the EM data. M.T., D.N., and I.P. processed and analyzed the EM data. H.S. provided guidance on EM data collection, processing, and evaluation. S.D., L.A.A, and I.P. performed the cellular assays and analyzed the cellular data with guidance from A.S. L.A.A., I.P. and S.G. wrote the manuscript.

## Supplemental information

Figures S1-S7, Tables S1, S2.

