## Supplementary material for "Arrestin recognizes GPCRs independently of the receptor state": Figure S1

### Supporting Information

*Address correspondence to:
Stephan Grzesiek
Focal Area Structural Biology and Biophysics, Biozentrum
University of Basel, CH-4056 Basel, Switzerland
Phone: ++41 61 267 2100
FAX: ++41 61 267 2109


Layara Abiko
Focal Area Structural Biology and Biophysics, Biozentrum
University of Basel, CH-4056 Basel, Switzerland
Phone: ++41 61 267 2100
FAX: ++41 61 267 2109


#### Supporting Tables

**Table S1.** Survey of published G protein, arrestin and GRK signaling strengths[a] of the β1AR and β2AR for selected orthosteric ligands and apo receptors.

| **Ligand/receptor** | **G protein** | **arrestin**[b] | **GRK phosphorylation**[c] | **Ref.** |
|---|---|---|---|---|
| apo β2AR-V2Rpp | 0 | 0 | 0.1 | (*1*)[d] |
| isoprenaline•β2AR-V2Rpp | 1 | 1 | 1 | (*1*) |
| carvedilol•β2AR-V2Rpp | 0.03 | 0.2 | 0.3 | (*1*) |
| apo β2AR$^{TYY}$-V2Rpp[e] | - | 0.07 | - | (*1*) |
| isoprenaline•β2AR$^{TYY}$-V2Rpp | - | 1 | - | (*1*) |
| carvedilol•β2AR$^{TYY}$-V2Rpp | - | 0.3 | - | (*1*) |
| apo β1AR | 0 | 0.2 | 0.4 | (*2*)[d] |
| isoprenaline•β1AR | 1 | 1 | 1 | (*2*) |
| carvedilol•β1AR | 0.05 | 0.6 | 0.7 | (*2*) |
| isoprenaline•β1AR | 1 | - | - | (*3*)[f] |
| formoterol•β1AR | 0.9-1.2 | - | - | (*3*) |
| cyanopindolol•β1AR | 0-0.39 | - | - | (*3*) |
| carvedilol•β1AR | 0-0.12 | - | - | (*3*) |

[a] Signaling strengths are given relative to the isoprenaline-induced response.
[b] quantified as receptor internalization.
[c] quantitative Western blot of receptor phosphorylation by GRK.
[d] For references Wisler *et al.* (2007) (*1*) and Kim *et al.* (2008) (*2*), the values were extracted from bar plots in the respective figures.
[e] β2AR$^{TYY}$ is the G protein-inactive mutant T68$^{2x39}$F,Y132$^{3x51}$G,Y219$^{5x58}$A.
[f] Reference Baker *et al.* (2011) (*3*) comprises various thermostabilized β1AR mutants.

**Table S2.** Cryo-EM data collection and image processing of carvedilol•$\beta_1$AR•arrestin2•V2Rpp complex.

| Data acquisition | |
|---|---|
| Instrument | Titan Krios G4 |
| Electron gun | Cold-FEG |
| Voltage (kV) | 300 |
| Energy filter | SelectrisX |
| Energy filter slit (eV) | 10 |
| Detector | Falcon 4i |
| Nominal Magnification | 120 kx |
| Pixel size (Å) | 0.658 |
| Electron fluency (e-/Å2) | 60 |
| Defocus range (μM) | 0.5 to 2.5 |
| Movies | 27,170 |
| **Data processing** | |
| Initial particle images (no.) | 3,235,599 |
| Final particle images (no.) | 38, 975 |
| Map resolution FSC 0.143 (Å) | N/A |
| EMPIAR codes for raw data | EMPIAR-12139 |
| EMDB codes for 3D maps | EMD-50623 |

#### Supporting Figures

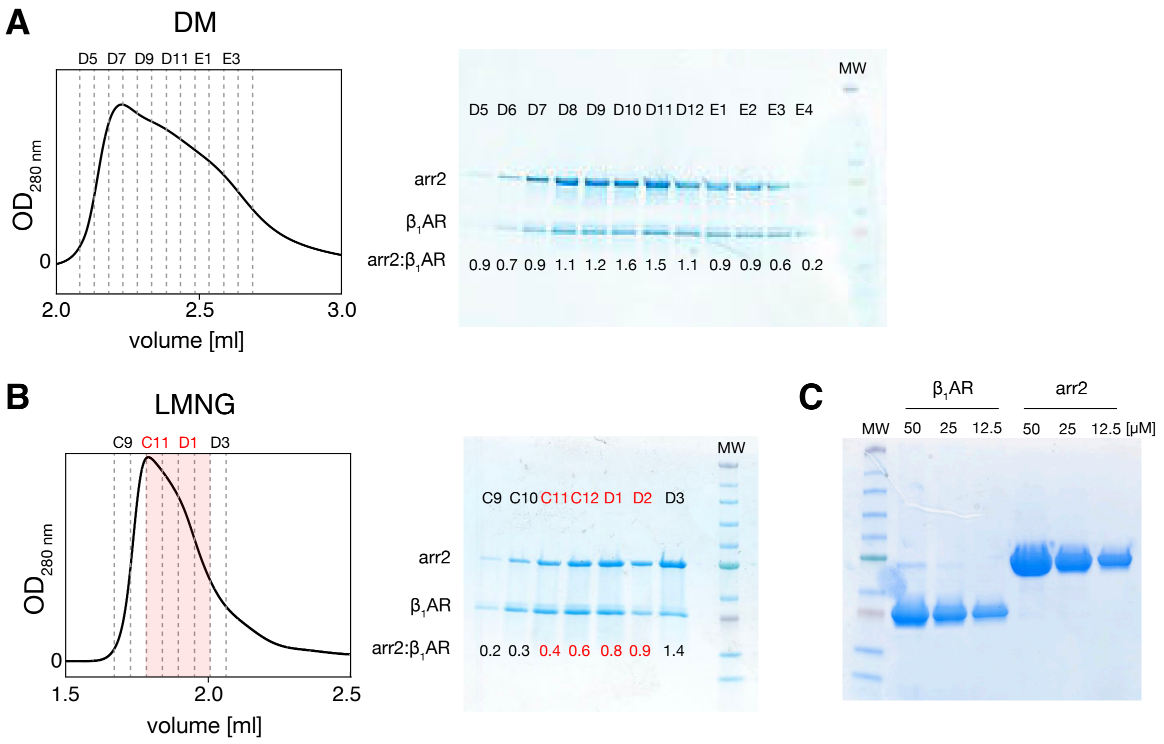


**Fig. S1: Carvedilol•β1AR•arrestin2•V2Rpp complex formation assessed by SEC and SDS-PAGE. (A)** in DM and **(B)** in LMNG micelles. SEC fractions are labeled at the top of the chromatograms. The fractions of the LMNG preparation used for cryo-EM grid preparation are highlighted in red. The bands in the individual fractions show that arrestin2 and β1AR are the only detected proteins. **(C)** Calibration of the Coomassie staining of β1AR and arrestin2 in a single gel for the indicated protein concentrations. Bands in all gels were quantified by ImageJ (*4*) and the molar ratios of arrestin2 and β1AR (arr2:β1AR) were calculated for all lanes in panels **(A)** and **(B)** using the calibration from panel **(C)**.

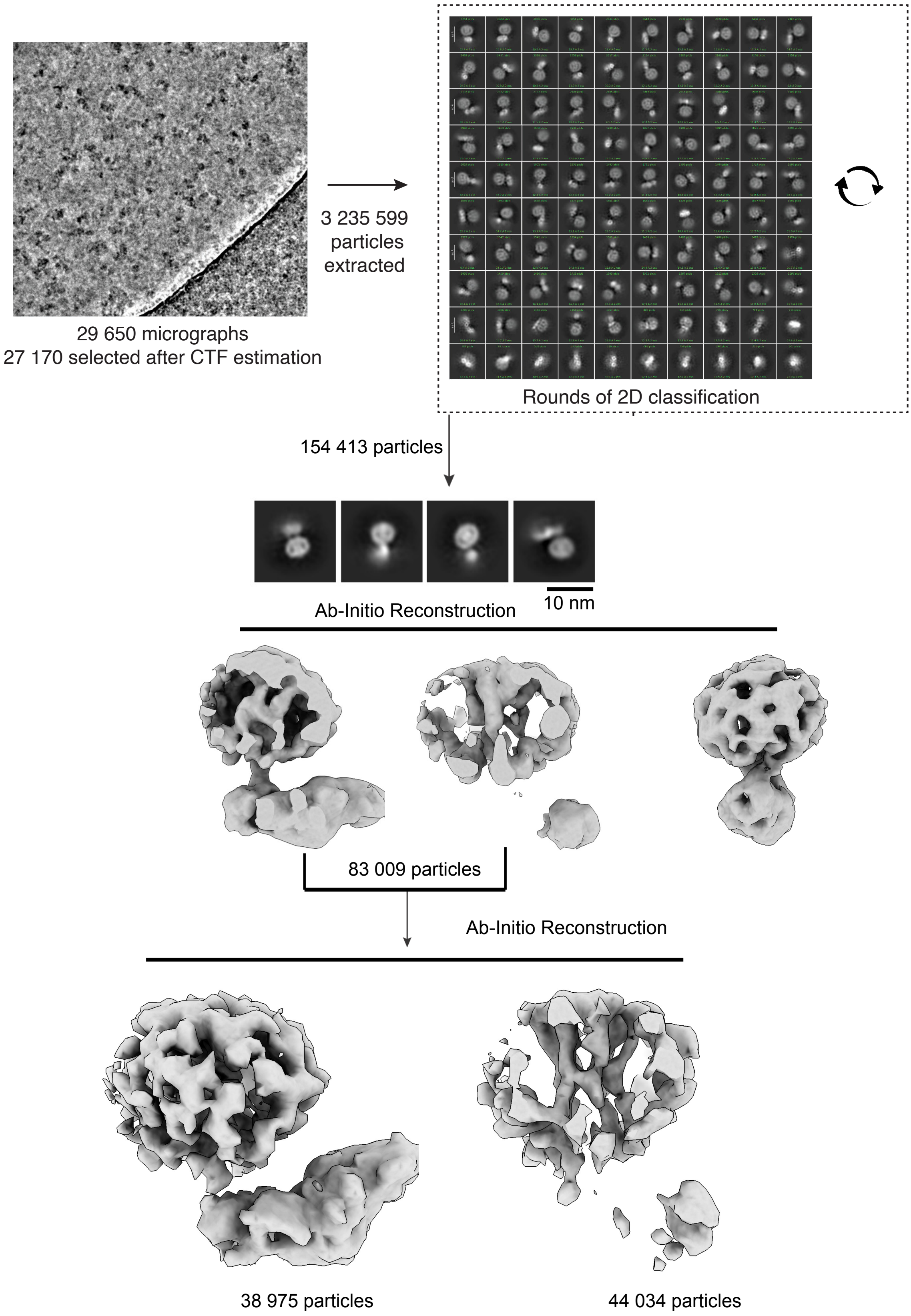


**Fig. S2: Cryo-EM data analysis workflow.** Cryo-EM processing summary of carvedilol•$\beta_1$AR•arrestin2•V2Rpp complex.

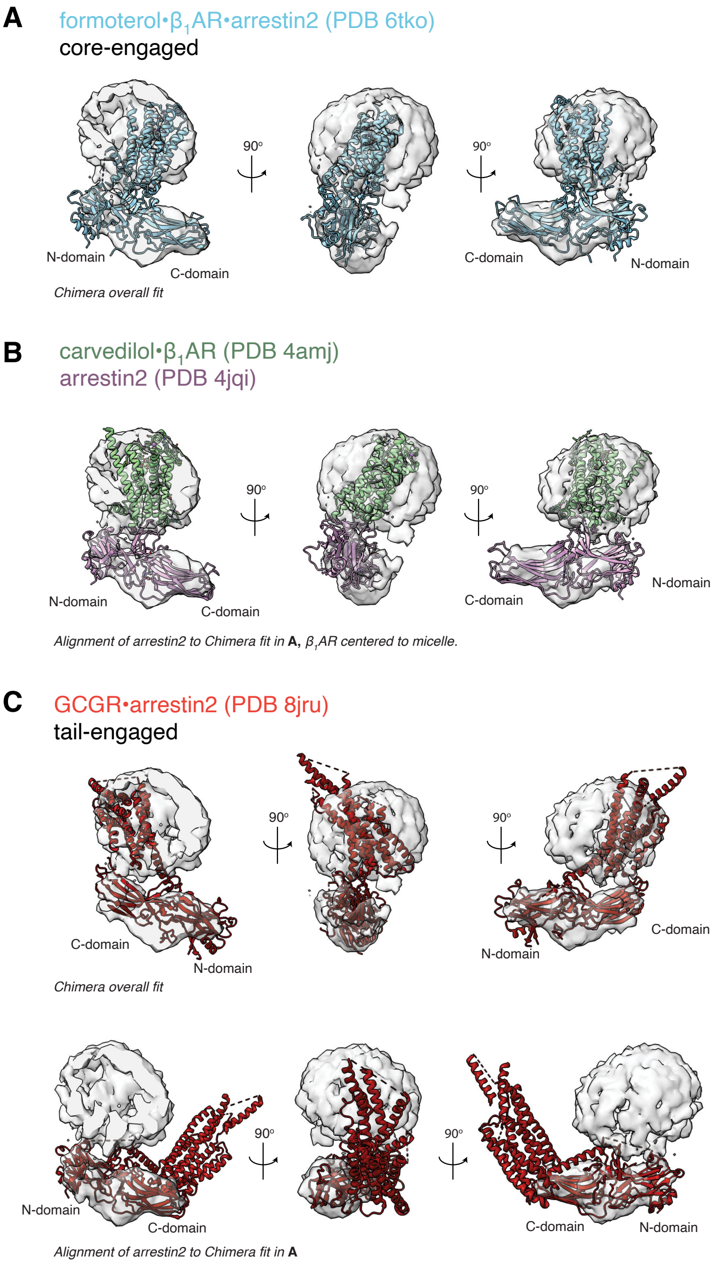


**Fig. S3: Comparison of the carvedilol•β1AR•arrestin2•V2Rpp complex cryoEM density to solved GPCR core- and tail-engaged arrestin complexes. (A)** Fit of the core-engaged formoterol•β1AR-V2Rpp•arrestin2 complex (PDB 6tko) to the cryoEM density using the 'Fit in Map' option of Chimera X. **(B)** as **(A)** but arrestin2 replaced by the arrestin2•V2Rpp (PDB 4jqi) and β1AR replaced by the β1AR•carvedilol (PDB 4amj) complex structures, with β1AR rotated by visual inspection to the center of the micelle density. **(C, top row)** Fit of the tail-engaged GCGR•arrestin2 structure (PDB 8rju) to the cryoEM density using the 'Fit in Map' option of Chimera X. **(C, bottom row)** GCGR•arrestin2 structure (PDB 8rju) aligned by the coordinates of arrestin2 to the fitted arrestin2 position of the formoterol•β1AR-V2Rpp•arrestin2 complex in panel **(A).**

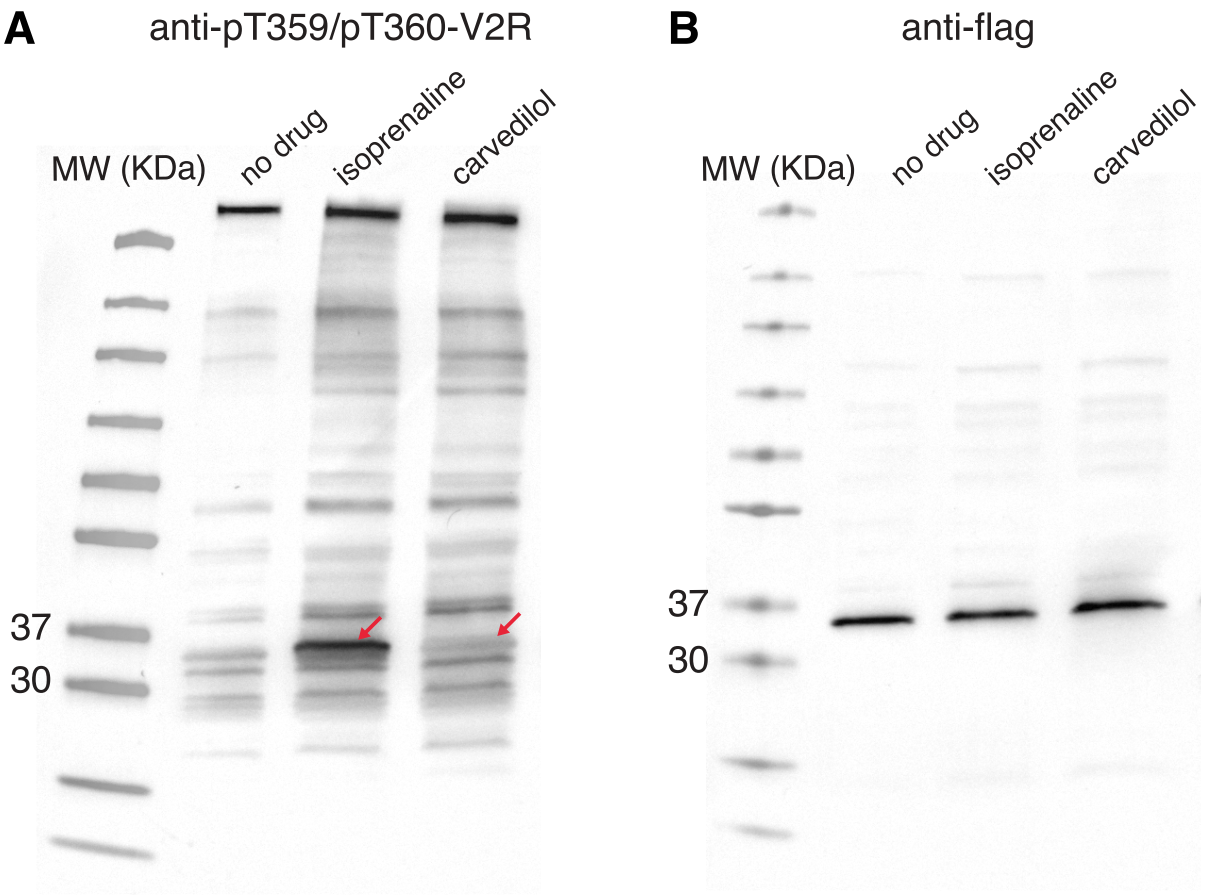


**Fig. S4: β₁AR phosphorylation in HEK-293 cells expressing β₁AR fused to V2Rpp.** Western blot images detected by a phosphosite-specific antibody (pT359/pT360-V2R) (**A**) or anti-flag tag **(B)**. Phosphorylation was stimulated by the addition of 10 μM of isoprenaline or carvedilol. No drug stimulation is used as a negative control. The bands for phosphorylated β₁AR-V2Rpp are indicated with red arrows. The intensity of the receptor bands in the anti-flag tag experiment was used to scale the phosphorylation obtained with anti-pT359/pT360-V2R relative to β₁AR-V2Rpp expression levels.

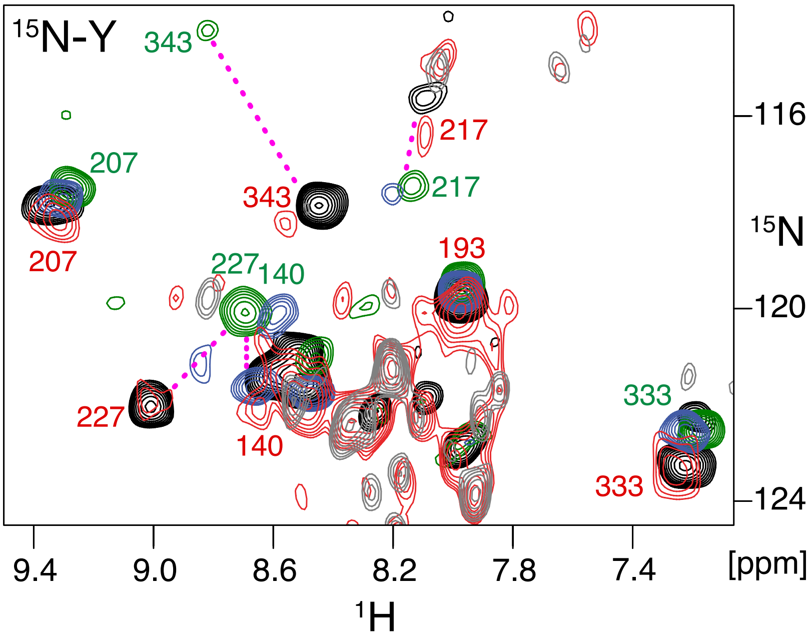


**Fig. S5: Determination of the β1AR conformation in various complexes by $^{1}$H-$^{15}$N tyrosine NMR.** Superposition of $^{1}$H-$^{15}$N TROSY spectra of $^{15}$N-tyrosine-labeled receptor in isoprenaline•β1AR$^{V129I}$•arrestin2•V2Rpp (red), isoprenaline•β1AR$^{V129I}$ (G protein-active, blue), isoprenaline•β1AR$^{V129I}$•Nb80 (G protein-active, green), isoprenaline•β1AR (G protein-preactive, black), and arrestin2•V2Rpp ($^{15}$N natural abundance, gray) complexes. Due to the low intensity of the $^{1}$H-$^{15}$N-tyrosine resonances from the isoprenaline•β1AR$^{V129I}$•arrestin2•V2Rpp complex (red), also $^{15}$N-natural-abundance resonances of free arrestin2 become visible in its spectrum. Resonances are marked with assignment information. Resonances connected by a dashed magenta line represent residues with two clearly distinguishable resonances for the G protein-inactive and -active conformations. No resonance corresponding to the G protein-active state (green) is observed in the carvedilol•β1AR$^{V129I}$•arrestin2•V2Rpp complex (red).

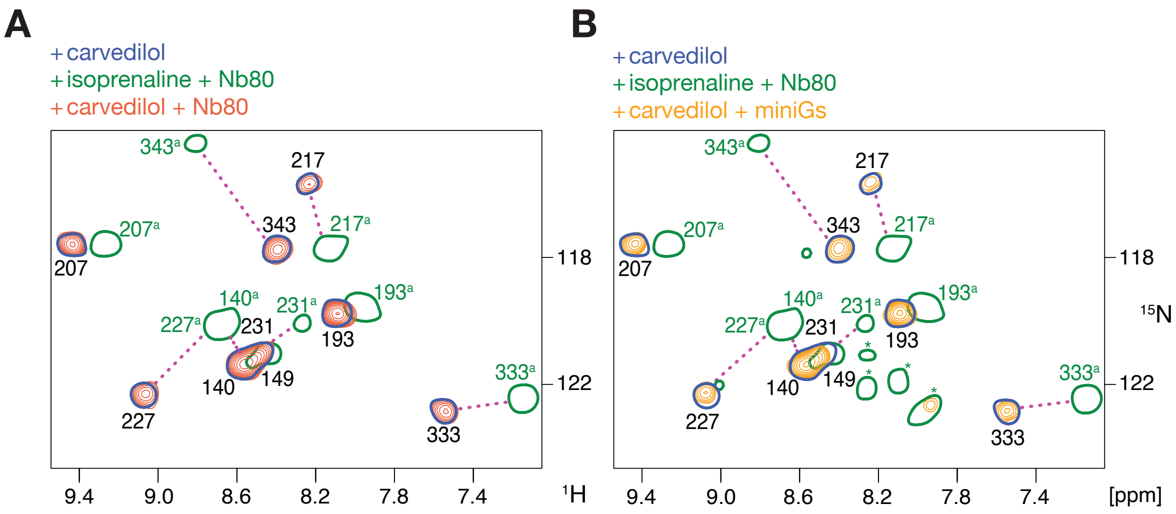


**Fig. S6: $^{1}$H-$^{15}$N-TROSY control experiment proving the absence of interactions between carvedilol•β$_1$AR and Nb80 or miniGs.** Comparison of $^{1}$H-$^{15}$N TROSY spectra of $^{15}$N-tyrosine-labelled carvedilol-bound β$_1$AR in the presence of Nb80 (**A**, orange) or miniGs (**B**, yellow) with those of carvedilol•β$_1$AR (blue) and isoprenaline•β$_1$AR•Nb80 (green) complexes as references for the G protein-inactive and -active states, respectively. For clarity, the resonances of carvedilol•β$_1$AR and isoprenaline•β$_1$AR•Nb80 are depicted using single contour lines. The resonances are labeled with assignment information and marked by an 'a' for the active conformation. Asterisks indicate spurious non-β$_1$AR resonances. Resonances of carvedilol•β$_1$AR + Nb80 (orange) and carvedilol•β$_1$AR + miniG$_S$ (yellow) spectra have the same chemical shifts and intensity as the carvedilol•β$_1$AR spectrum (blue), indicating no interaction.

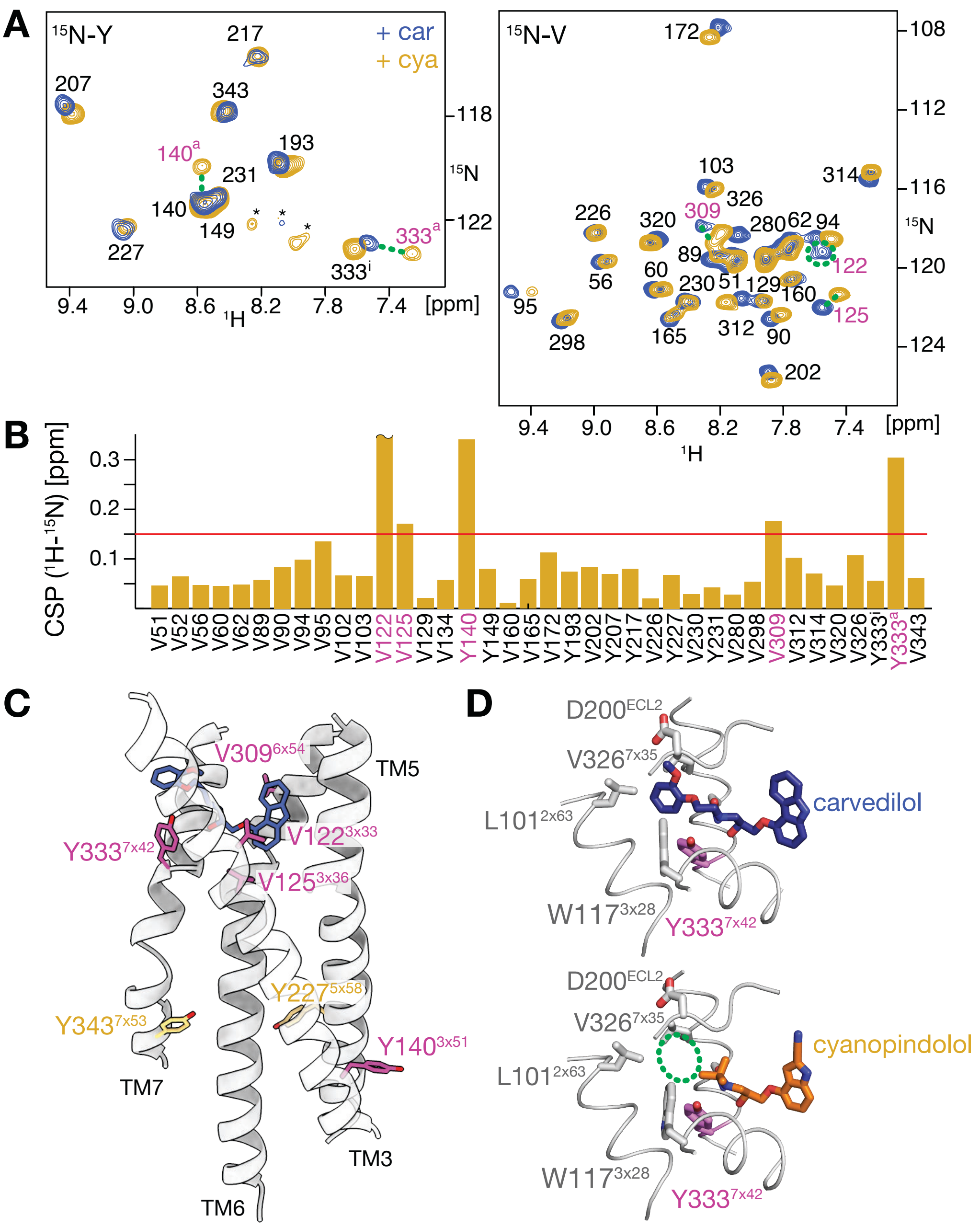


**Fig. S7: Comparison of cyanopindolol- and carvedilol-bound β1AR.** **(A)** Superposition of $^{1}$H-$^{15}$N TROSY spectra of $^{15}$N-tyrosine- (left) or $^{15}$N-valine-labelled (right) $\beta_1$AR in complex with cyanopindolol (orange) or carvedilol (blue). Resonances are marked with assignment information. Resonances connected by a dashed green line indicate residues which display large chemical shift difference between the cyanopindolol- and carvedilol-bound receptor. A dashed green circle indicates the disappearance of the V122 resonance in the cyanopindolol•$\beta_1$AR. Y333 exhibits two clearly distinguishable resonances corresponding to the inactive (‘i’) and active (‘a’) conformations. Resonances that are not assigned to any receptor residue are marked with ‘*’. **(B)** $^{1}$H-$^{15}$N chemical shift perturbation between cyanopindolol- and carvedilol-bound $\beta_1$AR. The red line represents a threshold of 0.15 ppm used to distinguish the most significant differences. The CSP bar for V122 capped at 0.35 ppm indicates that the resonance was bleached out due to exchange in the cyanopindolol-bound receptor. (**C)** The residues exhibiting significant chemical shift differences are shown as magenta sticks in the carvedilol•$\beta_1$AR crystal structure (PDB 4AMJ). Most of these residues are localized around the orthosteric binding site, with the exception of Y140$^{3x51}$, which is located on the intracellular side of the receptor. This perturbation is propagated through V122$^{3x33}$ and V125$^{3x36}$, which are in proximity to the ligand. The conserved tyrosines Y227$^{5x58}$ and Y343$^{7x53}$ which are essential for G-protein binding are shown as yellow sticks. (**D)** Close-up view of the $\beta_1$AR orthosteric binding site with carvedilol (top) and cyanopindolol (bottom). Residues of $\beta_1$AR in close proximity to the tail moiety of the ligands are labelled and shown as grey sticks. Y333$^{7x42}$, which exhibits a single resonance in the carvedilol•$\beta_1$AR complex, but two distinct resonances in the cyanopindolol•$\beta_1$AR complex, is shown as magenta sticks. The dashed green circle indicates the vacant space in the cyanopindolol•$\beta_1$AR complex, which is occupied by the anisole group of carvedilol, helping to stabilize TM7 in the carvedilol•$\beta_1$AR complex.
